# Distinct functions of Nup93 paralogs in tumor growth and Polycomb-mediated repression of JAK/STAT signaling

**DOI:** 10.64898/2026.08.28.747911

**Authors:** M. O’Sullivan, J. Hartmann, M. McLellan, D. Thuerauf, K. Bojorquez, G. Ulukaya, D. Hasson, P. Rangan, M. Capelson

## Abstract

Nuclear pore complexes (NPCs) are nuclear envelope (NE)-embedded protein assemblies that mediate nucleocytoplasmic exchange and interact with the genome, including binding of an NPC component Nup93 to Polycomb chromatin domains. Here, we investigated the *in vivo* relevance of this relationship in Drosophila, which unusually contains two distinct paralogs of Nup93. Interestingly, we identified a Nup93-2-specific tumorigenic phenotype in larval wings, where depletion of Nup93-2, but not Nup93-1, led to tumor-like overgrowth, reminiscent of Polycomb mutations. Consistently, our transcriptomic analysis revealed a wide-spread loss of gene silencing in Nup93-2-depleted wings, particularly in a Nup93-bound Polycomb domain spanning genes for activators of JAK/STAT signaling. Nup93 paralogs were not found to differ in their effect on NPC biogenesis but strikingly, showed differences in subnuclear localization patterns. While Nup93-1 co-localized exclusively with fully assembled NPCs, Nup93-2 exhibited only partial co-localization and was found at additional NE locations in a tissue-specific manner. Together, our results identify an *in vivo* silencing role of a Nup93 paralog and suggest that Nup93-2 may form a unique NE-associated complex that targets a subset of Polycomb domains containing growth-promoting genes.

## INTRODUCTION

Nuclear pore complexes (NPCs) are large multi-subunit complexes that span the nuclear envelope (NE) and function as selectively permeable channels for nucleocytoplasmic transport. NPCs are composed of approximately 30 distinct components termed Nucleoporins (Nups), which are arranged in distinct sub-complexes and structural features. The core NE-embedded structure of the NPC is an octameric ring, formed by repeated units of the Nup107-Nup160 outer-ring and the Nup93-Nup155 inner-ring sub-complexes [1,2]. This core scaffold interacts with Nups that make up the critical auxiliary structures such as the nuclear basket, the cytoplasmic fibrils and the inner channel. Core and auxiliary Nups vary in their stability of NPC association and related protein lifespan [3,4]. Whereas core Nups of the outer- and inner-ring sub-complexes are stably associated with the NPC and exhibit unusually long protein stability (and are thus termed stable Nups), auxiliary Nups show dynamic association with the NE-embedded NPC and higher protein turnover rates (and are thus termed dynamic Nups) [5–7].

Although highly conserved across eukaryotes and present in all cell types, NPC components have been linked to a variety of cell type-specific phenotypes and human pathologies, including cardiovascular and neurodegenerative conditions [8–11]. One of the most prominent disease connections for the NPC is the demonstrated roles of Nups in cancer [12,13]. Aberrant gene fusions of several Nups, such as Nup98, Nup214, and Tpr, are known to underlie several types of hematopoietic and gastric cancers [14–16], while the reduction of NPC levels or functionality was found to selectively inhibit oncogenic cell proliferation [13]. Mutations in specific Nups, both stable and dynamic, have also been shown to disrupt differentiation of specific cell lineages[12,17]. For example, the outer-ring sub-complex component Seh1 is involved in neural differentiation [18], while transmembrane component Nup210 is required for differentiation of muscle and immune cells [19,20] and dynamic Nup98 has been found to regulate hematopoietic development [21,22]. The mechanisms of many of these conditions remain incompletely understood, prompting the need to further investigate physiological and developmental roles of individual NPC components.

In addition to their canonical roles in nucleocytoplasmic transport, NPC components have been shown to interact with the genome and to influence expression and chromatin state of their gene targets across species [23,24]. The chromatin-binding functions of Nups have been found to regulate gene expression in developmental contexts and to control tissue-specific differentiation [25–28], highlighting a key mechanism via which Nup mutations can influence cellular physiology. Although commonly found at active genes [29–31], select Nups can also interact with heterochromatin and silent genomic regions [32–34] as well as with architectural features of the genome [26,35]. Understanding chromatin-binding roles of NPC components has been complicated by the intranuclear presence of dynamic Nups, which have been shown to primarily target actively transcribing or poised genes in the nuclear interior [29,30,36]. To distinguish the on-pore versus off-pore modes of Nup-genome contacts, chromatin mapping of stable NPC components has been used to determine genomic regions that are targeted preferentially to the NE-embedded NPCs [37]. Such analysis in Drosophila found that while the outer-ring component Nup107 is found primarily at active chromatin states, binding of the inner-ring sub-complex component Nup93 is enriched for genomic regions repressed by Polycomb group (PcG) proteins. This binding was found to be functionally important, as both optimal gene silencing and long-range clustering of Polycomb domains were shown to be dependent on Nup93 in S2 culture cells [37]. These findings revealed that NE-embedded Nups can function as scaffolds for Polycomb-repressed heterochromatin domains, which impacts the critical roles that Polycomb machinery plays in cell differentiation and epigenetic memory.

The identified role of Nup93 in the maintenance of repressed chromatin domains is supported by findings in mammalian cells [38–40] and by multiple studies that demonstrated a relationship between heterochromatin and Nups of the same inner-ring sub-complex. In fission yeast, the Nup93 homolog and members of its sub-complex support the maintenance and nuclear clustering of constitutive heterochromatin [41]. Another defining component of this sub-complex, Nup155 (Nup154 in Drosophila) binds and facilitates sub-telomeric heterochromatin in budding yeast [34], interacts with Histone Deacetylases (HDACs) in mammalian cardiomyocytes [42], and promotes establishment of heterochromatin in the Drosophila germline [33]. Together, these findings indicate that inner-ring Nups carry an evolutionarily conserved role in nuclear organization and maintenance of heterochromatic regions. Yet the precise mechanisms of this role remain poorly defined, and the developmental significance of the NPC-Polycomb relationship is still unclear.

Homologs of Nup93 are present across eukaryotes, and as an inner-ring component, it is structurally required for nuclear pore biogenesis and assembly [43]. Within the NPC, Nup93 and the inner-ring sub-complex are responsible for interactions with inner channel Nups, which create the selective permeability barrier via their extended repeats of Phenylalanine and Glycine (FG repeats) [44–46]. Nup93 is also one of the most stable Nups within the NPC [47], which may make its cellular function particularly consequential. Mutations and expression changes in Nup93 are linked to several human diseases, which, once again, are quite cell type-specific.

Mutations in Nup93 have been shown to cause nephrotic syndrome [48,49] and congenital ataxia [50], while elevated expression and mutations of Nup93 have been linked to oncogenesis and cancer aggressiveness [51,52], at least partly via aberrant activation of Wnt signaling [53]. Whether the chromatin-binding role of Nup93 contributes to these conditions is unknown. In addition to Polycomb domains, Nup93, together with other NPC components, has been shown to target a subset of super-enhancers and to promote cell type-specific expression in mammalian cells [54,55]. Thus, understanding physiological roles of Nup93 and its function in gene regulation will inform mechanisms of cell identity and human disease linked to NPCs.

To gain further understanding of these roles and to begin probing the developmental relevance of the discovered NPC-Polycomb relationship, we investigated the phenotypic consequences of depleting Nup93 in Drosophila tissues. Interestingly, Drosophila possesses two distinct paralogs of Nup93, Nup93-1 and Nup93-2, the genes for which are carried on different chromosomes. The presence of two versions of Nup93 is unique among model organisms and suggests that Nup93 paralogs may have evolved distinct functions or separated an existing set of functions between individual paralogs. Our phenotypic and transcriptomics analysis revealed distinct tissue-specific requirements and gene targets of the two paralogs, and significantly, identified a tumorigenic and gene repressive phenotype specific to loss of Nup93-2. Unexpectedly, we observed that in addition to normal NPC localization, Nup93-2, but not Nup93-1, may be localized to unique non-NPC locations within the NE. Together, our findings reveal that Nup93 paralogs regulate distinct developmental processes and support a model, where Nup93-2 is part of a unique NE-embedded complex that helps silence a subset of Polycomb-repressed growth-promoting genes.

## RESULTS

### 1. Drosophila Nup93 paralogs exhibit unique requirements in tissue-specific development

Drosophila Nup93 paralogs, Nup93-1 and Nup93-2, share 45% amino acid identity and 65% similarity, and carry a similar level of homology to the human Nup93 (Figure 1A, S1A). Our original ChIP-seq studies were carried out with an antibody that recognizes a domain with significant similarity between the paralogs, thus it is likely that the reported chromatin binding of Nup93 [37] represents binding of both paralogs. To determine whether Nup93 paralogs play distinct roles in the organism and whether a specific Nup93 paralog is preferentially involved in gene silencing, we depleted Nup93-1 and Nup93-2 individually in different tissue types using GAL4/UAS-driven RNAi. Interestingly, we found that while whole-body RNAi knock down (KD) of either paralog lead to early (larval) lethality, tissue-specific KD produced unique lethality outcomes (Figure 1B). For example, Nup93-1 RNAi resulted in larval and pupal lethality when driven by muscle-specific (Mef2-) and neuron-specific (Elav-) GAL4, respectively, while Nup93-2 RNAi produced semi-lethal or fully viable adults with the same drivers. In contrast, KDs in epithelial tissues of larval imaginal discs, using wing-specific Nub-GAL4 and eye-specific Ey-GAL4, showed a reverse scenario – RNAi of Nup93-2 was found to be pupal lethal with both drivers, while RNAi of Nup93-1 produced fully viable adults (Figure 1B). Since both Nup93 RNAi lines can produce lethality albeit in different tissues, insufficient level of knock down is unlikely to explain these phenotypic differences. Together, the obtained tissue-specific lethality pattern suggests that different cell types possess unique requirements for individual Nup93 paralogs.

**Figure 1.**
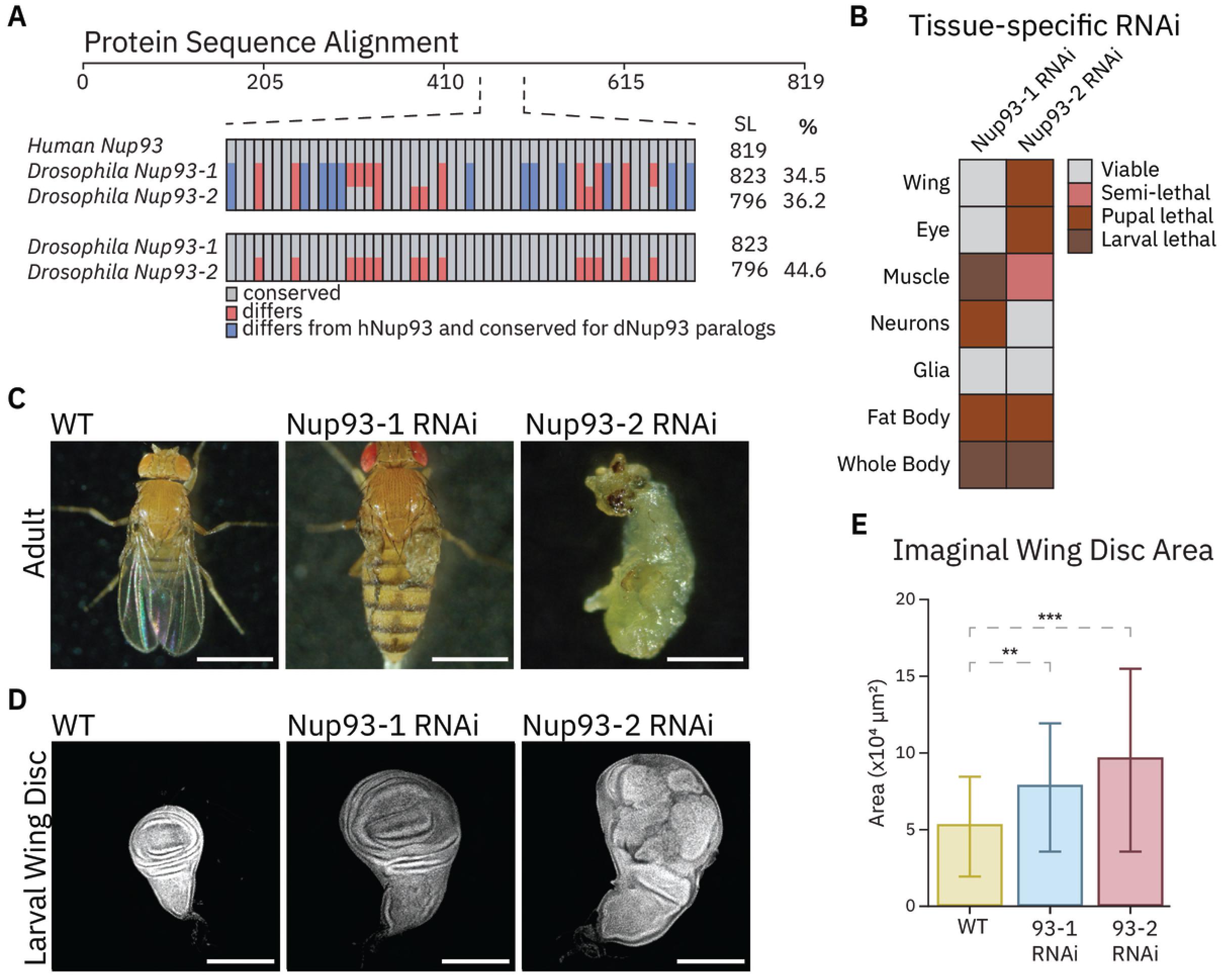
Drosophila Nup93-1 and Nup93-2 exhibit distinct tissue-specific phenotypes. (A) Protein Sequence Alignment: comparison of protein sequences of human Nup93 and Drosophila Nup93 (top alignment) and of just Drosophila Nup93 paralogs to each other (bottom alignment). Gray=conserved throughout the comparison, red=differs from one another, blue=differs from hNup93 and conserved for dNup93 paralogs. (B) Tissue-specific RNAi: UAS-RNAi against each dNup93 paralog was induced by different tissue-specific Gal4 drivers (see Methods) and resulting larval and adult lethality scored. Note the differences between the two paralogs in tissue-specific lethality. (C) Adult of pupal phenotypes of Nub-Gal4-driven RNAi against each dNup93 paralog compared to wild type. Nup93-2 RNAi is pupal lethal, while Nup93-1 RNAi is not (Brightfield microscopy, 1x objective, scale bar 1mm). (D) Wing imaginal discs of third instar larvae of Nub-Gal4-driven RNAi against each dNup93 paralog compared to wild type. Representing their tissue-specific differences. Nup93-1 RNAi shows increased size but normal morphological patterning, while Nup93-2 RNAi shows severe tumor-like disorganization and overgrowth (Confocal microscopy, 10x objective, scale bar 200µm). (E) Quantification of imaginal wing disc total area in genotypes from (D). One-way ANOVA + Dunnett’s statistical analysis in Prism (n=17-69 wings per genotype, ns=not significant, *p<0.05, **p<0.01, ***p<0.001).

The observed lethality of wing-specific Nup93-2 KD was particularly interesting, since generally flies do not require wings for viability. While Nub-GAL4-driven Nup93-1 KD produced malformed wings in otherwise normal adults, Nub-GAL4-driven Nup93-2 KD led to early pupal lethality (Figure 1C). To further understand this lethal phenotype, we examined wing imaginal discs, which are precursor structures to adult wings, in late third instar larvae of each Nup93 paralog Nub-GAL4-driven KD. We found that Nub-GAL4-driven Nup93-2 KD resulted in a striking tumorigenic phenotype in the wing disc, characterized by over-proliferation and drastic dis-organization of wing epithelial tissue (Figure 1D-E). Wing discs of Nub-GAL4-driven KD of Nup93-1 appeared larger yet showed normal morphology and no epithelial dis-organization.

The discovered tumor-like phenotype of Nup93-2 suggests a specific role of this paralog in regulation of growth signaling and tumor suppression. Generally, mutations that produce imaginal disc tumors in flies are known to cause larval and pupal lethality [56,57], which helps explain the observed Nup93-2 wing-specific lethality. Significantly, one class of mutations with this phenotype is PcG mutations, particularly those in components of the Polycomb Repressive Complex 1 (PRC1) [56,58,59]. The tumor-like phenotype of Nup93-2 appears to be uncommon among Nups as other NPC components we depleted with Nub-GAL4 did not produce wing tumors (Figure S1B), further suggesting that the uncovered tumor suppressor function of Nup93-2 may be related to its connection to the Polycomb pathway.

### 2. Transcriptomics of wing discs depleted for each Nup93 paralog reveal paralog-specific gene targets and widespread gene upregulation

To further investigate the observed phenotypic differences between Nup93 paralogs and the tumorigenic phenotype of the Nup93-2 KD in the wing, we carried out RNA sequencing (RNA-seq) analysis on wing discs. To this end, we isolated batches of wing discs from wandering third instar larvae of control (Nub-GAL4, also referred to as wild-type or WT), Nub-GAL4-driven Nup93-1 KD and Nub-GAL4-driven Nup93-2 KD genotypes, in triplicate for each, and performed mRNA-specific RNA-seq on isolated RNA (Figures 2, S2). Comparison of differentially expressed genes (DEGs) between Nup93-1 or Nup93-2 KD vs WT RNA-seq datasets demonstrated a predominant upregulation of gene expression for either paralog depletion (Figure 2A-C), with more genes upregulated than downregulated in either Nup93-1 or Nup93-2 KD. These findings support the previously identified functional role of Nup93 paralogs in gene silencing [37] and demonstrate it in a developing organism. Importantly, each paralog was targeted specifically by its RNAi, such that Nub-GAL4-driven Nup93-1 KD exhibited normal levels of Nup93-2 and Nub-GAL4-driven Nup93-2 KD had normal levels of Nup93-1 (Figure 2D), indicating that these gene expression changes are specific to each paralog.

**Figure 2.**
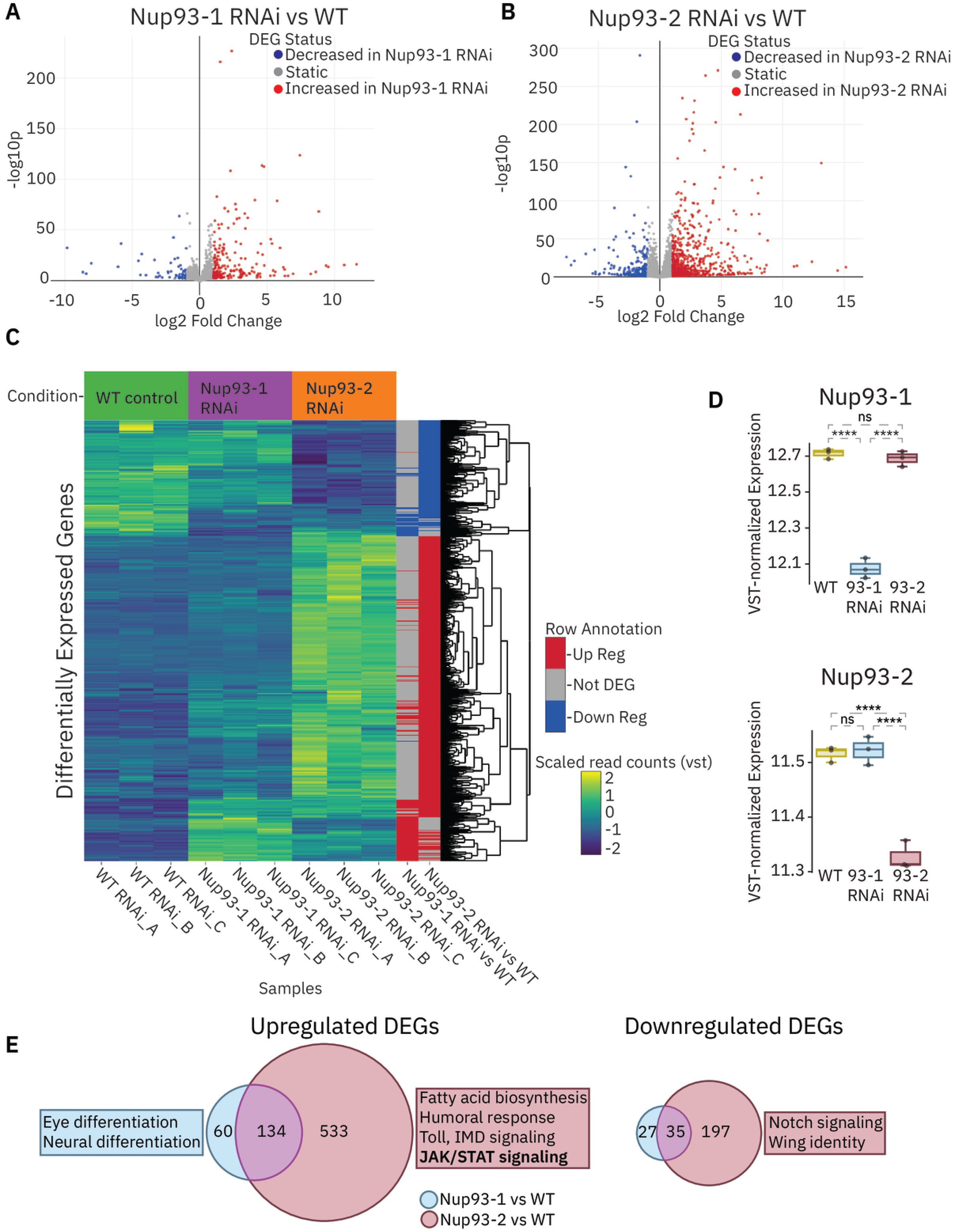
Wing-specific depletions of Nup93 paralogs lead to widespread gene upregulation and reveal paralog-specific gene targets. (A) Results of RNA-sequencing of larval wing discs from Nub-Gal4-driven RNAi of individual Nup93 paralogs vs controls – volcano plot representing gene expression changes in Nup93-1 RNAi vs control (control genotype - Nub-Gal4 crossed to *w^1118^*). (Filtered by p<0.05, log2FoldChange >1). (B) Volcano plot representing gene expression changes in Nup93-2 RNAi vs control (Filtered by p<0.05, log2FoldChange >1).(C) Heatmap for cluster analysis of differentially expressed genes (DEGs) between Nub-Gal4-driven RNAi of individual Nup93 paralogs vs controls, as indicated. Each genotype has 3 replicates. Note the large unique category of upregulated genes in Nup93-2 RNAi. (D) Variance stabilized (VST) normalized expression of Nup93-1 and Nup93-2 from the RNA-seq datasets, demonstrating specific knockdown of each paralog in the appropriate RNAi condition. (Wald tests adjusted using Benjamin-Hochberg false discovery rate (FDR) analysis performed in R using DESeq2 package. n=3, ns=not significant, ****p<0.0001). (E) Overlap between upregulated DEGs of Nup93-1 vs Nup93-2 RNAi or downregulated DEGs of Nup93-1 vs Nup93-2 RNAi. Nup93-2 RNAi DEGs shown as red, Nup93-1 RNAi DEGs shown as blue. Enriched functional categories from non-overlapping genes obtained with FlyEnricher (see Methods).

Gene upregulation was particularly prevalent for Nup93-2 KD, with 667 genes upregulated more than 2-fold (232 down-regulated), compared to 194 upregulated in Nup93-1 KD (62 down-regulated). Comparison of the 3 datasets showed that the largest category of DEGs is genes that are strongly de-repressed in Nup93-2 KD (Figure 2C). Interestingly, some of the genes upregulated in the Nup93-2 KD were also upregulated in the Nup93-1 KD but to a considerably smaller degree (Figure 2C-E). Around 70% of upregulated DEGs and 55% of downregulated DEGs in Nup93-1-depleted wings were shared with DEGs in Nup93-2-depleted wings. On the other hand, 80% of DEGs upregulated in Nup93-2-depleted wings were unique to Nup93-2 (Figure 2C-E), suggesting that in the developing wing, Nup93-2 plays a particularly prevalent gene silencing role. This difference is unlikely to be due to differences in the levels of RNAi depletion, since the depletion levels of the two paralogs appear comparable (Figure 2D) and the same RNAi line of Nup93-1 is highly lethal in muscle and neural tissues (Figure 1B).

Enrichment analysis of paralog-specific DEGs using FlyEnrichr [60] revealed several signaling pathways among unique Nup93-2-KD-upregulated DEGs, including innate immunity pathways such as IMD and Toll, and significantly, the cell proliferation- and cancer-associated JAK/STAT signaling [61,62] (Figure 2E). Interestingly, among Nup93-1-KD-upregulated DEGs, enriched pathways included genes involved in neural differentiation (Figure 2E), which is consistent with the sensitivity of developing neurons to the KD of this paralog (Figure 1B). The cancer-related gene expression changes associated with Nup93-2 highlighted the potential mechanism of the tumor-like wing phenotype of Nup93-2 KD, prompting us to focus on the JAK/STAT signaling pathway.

### 3. Genes for JAK/STAT ligand Unpaired are direct targets of Nup93 and Polycomb-mediated repression

Persistent activation of JAK/STAT signaling has been widely implicated in human carcinogenesis [63] and is known to cause or commonly occur in Drosophila tumors [62,64]. The JAK/STAT pathway is hyperactivated in imaginal disc tumors caused by PcG mutations, and the JAK/STAT-activating cytokine ligands encoded by Unpaired (Upd) genes Upd1, Upd2 and Upd3 are known to be direct targets of Polycomb-mediated silencing [56,65,66]. Upd genes become highly derepressed in PRC1 mutants, and inhibiting JAK/STAT activation has been shown to rescue the tumor phenotypes of PRC1 mutations [56,65], demonstrating that JAK/STAT signaling is a key driver of Polycomb-based tumorigenesis.

Similarly to these effects of PRC1 mutations, we found that all three Upd genes, which are present as a gene cluster and a Polycomb domain in wing discs [67,68], became strongly derepressed in the Nup93-2 wing KD (Figure 3A, 3C). Importantly, the gene expression changes of Nup93-2 and Nup93-1 depletions could stem from either transport-related or chromatin-binding functions of Nup93. To help distinguish these possibilities, we analyzed our previously determined pan-Nup93 chromatin binding relative to the Upd gene cluster and observed robust ChIP-seq peaks of Nup93 at each of the Upd genes (Figure 3A). Although the Nup93 ChIP-seq dataset was obtained from S2 cells, the co-localization of Nup93 with wing-specific ChIP-seq of Polycomb (Pc) and areas of H3K27Me3 enrichment (Figure 3A-B) parallels the reported association of Nup93 with Polycomb domains in S2 cells [37]. Interestingly, we found that additional and seemingly unrelated genes located within the Upd cluster, such as CG15059 (Figure 3A, 3C, Figure 3SA) are also upregulated in the Nup93-2 KD, further supporting the idea that this de-repression is a chromatin-based mechanism and stems from overall loss of silencing around Nup93 binding sites.

**Figure 3.**
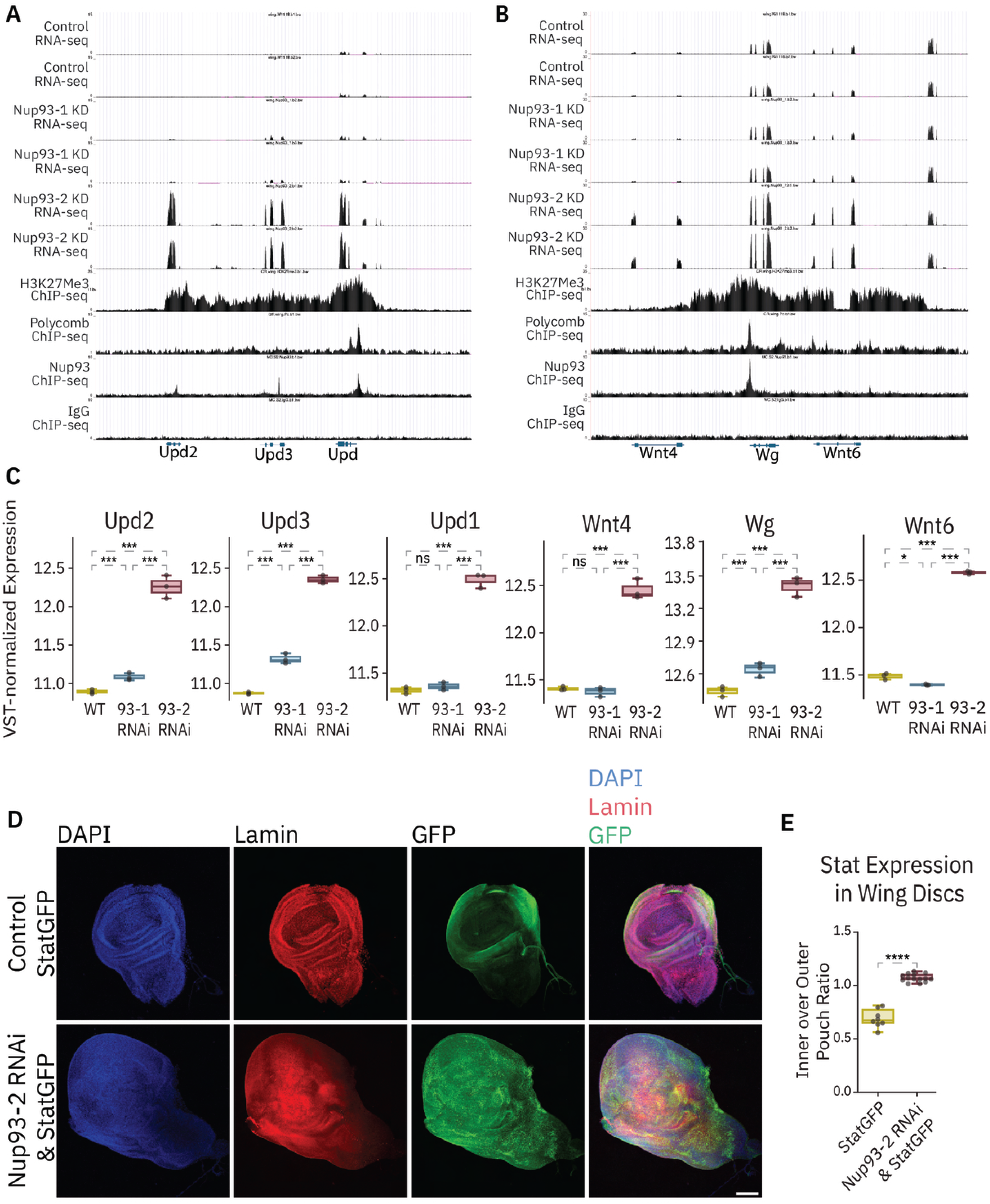
Loss of Nup93-2 results in de-repression of select Polycomb domains and aberrant activation of JAK/STAT signaling. (A) Comparison of a Polycomb domain and chromatin binding of pan-Nup93 at the Upd1/2/3 locus to RNA-seq datasets from Nup93-1 and Nup93-2 knockdowns (wing discs), using published ChIP-seq datasets of Pc and H3K27Me3 (wing discs, GSM3424769 and GSM3424770, from GSE121028) and of pan-Nup93 (S2 cells, GSE135610). (B) Comparison of a Polycomb domain and chromatin binding of pan-Nup93 at the Wg/Wnt locus to RNA-seq datasets from Nup93-1 and Nup93-2 knockdowns RNA-seq analysis, as in (A). (C) Normalized expression of Upd and Wg/Wnt genes, derived from RNA-seq datasets. (Wald tests adjusted using Benjamin-Hochberg false discovery rate (FDR) analysis performed in R using DESeq2 package. n=3, ns=not significant, *p<0.05, ***p<0.001). (D) Immunofluorescent staining in wild-type and Nup93-2-depleted wing discs for STAT-GFP expression (20x objective, scale bar 100µm). Tissue was stained with DAPI (blue), Lamin (red) for NE/nuclear lamina, and GFP (green) for Stat. (E) Stat-GFP expression, as in (D), was quantified as a ratio of the inner vs the outer region of the wing pouch in control vs Nup93-2 KD larvae. A one-way ANOVA + Dunnett’s statistical analysis was done in Prism (n=10).

We observed a similar behavior at another gene cluster linked to growth control and tumorigenesis – the Wingless (Wg) gene and nearby related Wnt genes Wnt4 and Wnt6 (Figure 3B). Wg signaling and its mammalian counterpart, Wnt signaling, are also widely implicated in human and Drosophila tumorigenesis [69] [70]. Similarly to the Upd gene cluster, the Wg and Wnt6 genes are part of a large Polycomb domain that shows nearly identical chromatin binding of Nup93 and Pc (Figure 3B). The Wnt4 gene lies immediately adjacent to the Wg/Wnt6-spanning Polycomb domain and its regulatory elements may be under its control. All 3 genes are upregulated specifically in the Nup93-2 KD (Figure 3B-C). As observed at the Upd locus, intervening and neighboring genes in the Wg region, such as Ndae1 are also upregulated in the Nup93-2 KD (Figure S3A). The observed domain-wide de-repression of Wg/Wnt4/Wnt6 gene region once again suggests a chromatin-based mechanism for the gene silencing role of Nup93-2. Wg signaling upregulation is also consistent with decreased Notch signaling that we observed in downregulated DEGs of Nup93-2 KD (Figure 2D), since Wg and Notch pathways are known to have antagonistic effects on each other [71].

To validate the upregulation of these pathways in Nup93-2 wing KD, we assayed for JAK/STAT and Wg signaling via immunofluorescent staining (Figure 3D-E, S3B). To this end, we utilized a GFP reporter, which contains binding sites for the main transcription factor of JAK/STAT signaling, Stat92E (10XSTAT-GFP). Whereas 10XSTAT-GFP staining highlighted the normal JAK/STAT activity pattern and was absent in the pouch region of the wild-type wing disc, we observed high JAK/STAT activity throughout the wing disc in Nub-GAL4-driven Nup93-2 KD (Figure 3D-E). Similarly, we detected elevated Wg staining throughout the wing pouch area of Nup93-2-depleted wing discs, in contrast to the wild-type pattern of narrow Wg staining at the dorsal/ventral boundary (Figure S3B). These results confirm our RNA-seq analysis and support our hypothesis that Nup93-2 functions as a tumor suppressor through a chromatin-binding role in silencing of JAK/STAT and Wg genes.

### 4. Nup93-2 is required for repression of Upd genes independently of tissue context or cell viability

Epithelial tumors in Drosophila imaginal discs have also been shown to arise via activation of the JNK signaling pathway in response to damage or stress, which in turn can activate JAK/STAT signaling [72–74]. One possibility is that the tumorigenic phenotype of Nup93-2 KD may arise in a more indirect manner from compromised NPC assembly, which may lead to increased cell death, JNK activation and associated compensatory cell proliferation [75] [76]. Interestingly, a previous study found that combining depletion of another NPC component, the dual Nup98-Nup96 gene, with genetic inhibition of apoptosis also leads to a tumorigenic phenotype in the Drosophila larval wing disc [77]. This Nup98-Nup96 phenotype was found to exhibit elevated JNK signaling and hallmark features of apoptosis-induced compensatory proliferation [78]. To gain insight into direct roles of Nup93-2, we aimed to determine whether loss of Nup93-2, relative to Nup93-1, can preferentially de-repress Upd genes or cause cell death outside of the tissue context, or differentially impact NPC levels. Since the tumorigenic phenotype appears to be unique to the Nup93-2 paralog, determining its effects relative to the Nup93-1 paralog would shed light on the tumor-associated mechanisms of Nup93-2.

To address these questions, we depleted either Nup93-1 or Nup93-2 or both, as well as Nup107 or PRC1 component polyhomeotic-p (ph-p) or control White gene, in Drosophila S2 embryonic culture cells via RNAi, and tested expression of several PcG gene targets (Figures 4, S4). We found that similarly to wing tissues, KD of Nup93-2 had a unique effect on Upd gene expression, leading to a significant upregulation of Upd2 and Upd3 as measured by RT-qPCR (Figure 4A). This de-repressive effect was stronger than a mild but non-significant upregulation we observed upon Nup93-1 KD, and for Upd3, it recapitulated the effect of depleting ph-p.

**Figure 4.**
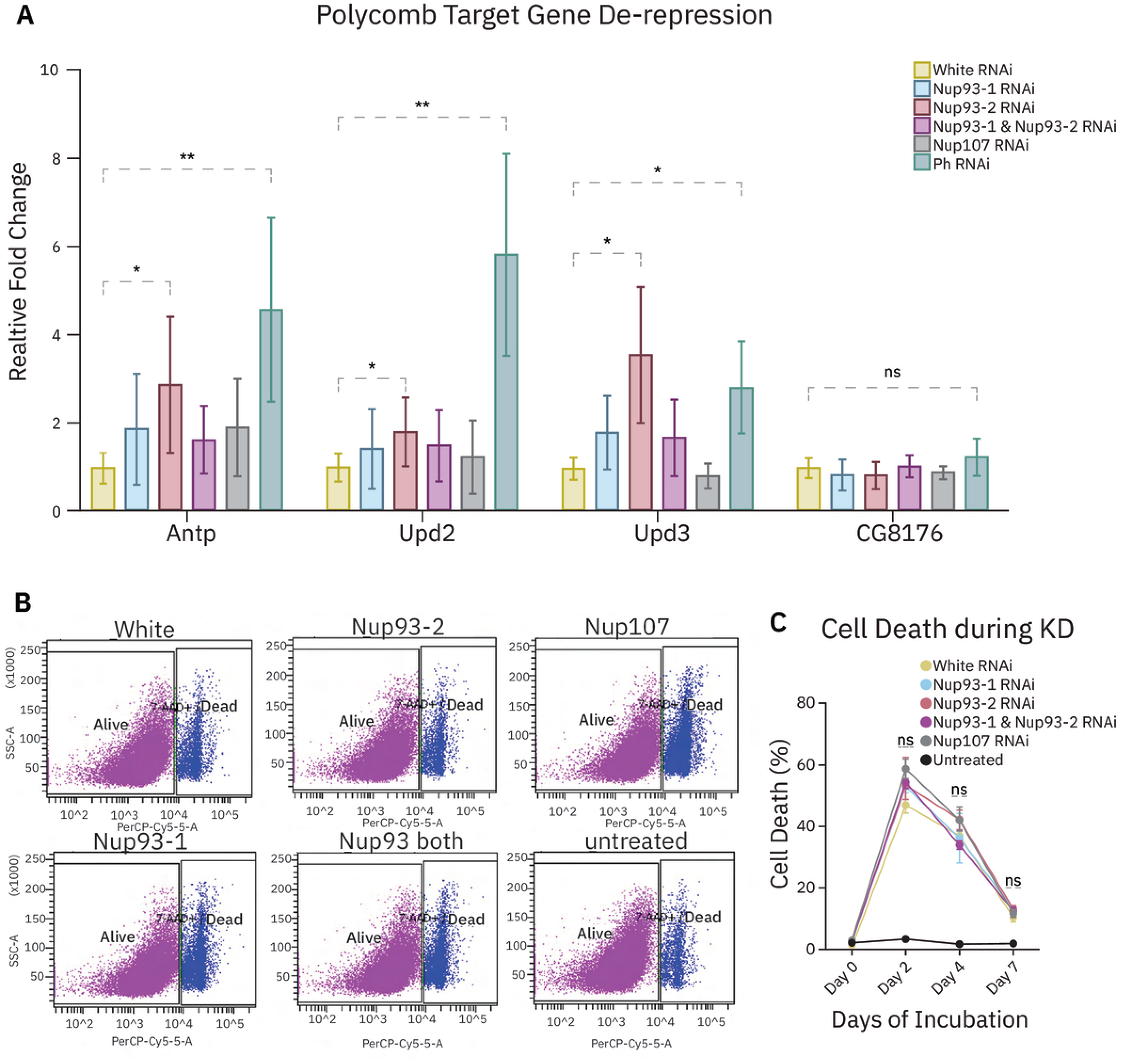
Nup93-2 functions in silencing of Upd genes across cell types and independently from effects on cell viability. (A) Normalized expression levels (relative fold change) of indicated Polycomb target genes and control CG8176 gene in S2 cells RNAi-treated for Nup93-1, Nup93-2, both Nup93 paralogs, Nup107, or positive control Ph-p or negative control, White, assessed via RT-qPCR, normalized to beta-tubulin expression. (one-way ANOVA with Holm-Sidak multiple comparisons test. n=3 biological replicates, *p<0.05, **p<0.01, ***p<0.001). (B) Flow cytometry viability analysis of S2 cells, RNAi-treated as in (A) and labeled appropriately, using 7-AAD stain. (C) Percentage of dead cells, calculated from flow cytometry analysis as in (B), obtained during a shown time course of RNAi knockdowns (KD) for indicated genes.

Nup93-2 KD also led to upregulation of a known Hox gene target of Polycomb silencing, Antp, paralleling the generally stronger silencing effect of Nup93-2 we observed in larval wing discs. Expectedly, RNAi against Nup107 did not lead to any upregulation of the tested PcG gene targets, and expression of a non-PcG control gene was not affected by any of the depletions (Figure 4A). Interestingly, depleting both Nup93-1 and Nup93-2 led to a weaker de-repression of any of the tested genes than KD of Nup93-2 alone. Although the reasons for this are presently unclear, it does not appear to stem from insufficient levels of Nup93-2 depletion (Figure S4).

In parallel to assessing gene expression, we also analyzed cell viability of the RNAi-treated S2 cells. To this end, we carried out flow cytometry to determine the fraction of dead cells during the time course of RNAi treatments for Nup93-1, Nup93-2, both Nup93 paralogs, Nup107, control White, or untreated cells (Figures 4B-C, S4B). Although we observed relatively high levels of general cell death upon all RNAi treatments relative to untreated cells, our analysis did not identify significant differences in dead cell counts between control dsRNA-treated cells and cells depleted for either Nup93 paralog (Figure 4B-C). We similarly did not observe significant differences in manual cells counts that we conducted for control, Nup93-2-depleted and Nup93-1-depleted cells over a similar RNAi treatment time course (Figure S4C), suggesting that upregulation of Upd genes that occurs in Nup93-2-depleted but not Nup93-1-depleted S2 cells is not explained by a differential effect on cell viability. Together, our findings in S2 cells indicate that Nup93-2 functions in repression of Upd genes across cell types and independently of cell death, supporting a direct mechanism in gene regulation.

### 5. Nup93 paralogs exert similar effects on NPC levels

To further investigate the tumorigenic phenotype of Nup93-2 and the possible functional differences between Nup93 paralogs, we considered several possibilities. One of these possibilities is that the Nup93 paralogs have unique functions in NPC assembly. In S2 cells, we were able to uncouple the de-repressive effect of Nup93-2 KD on Upd gene expression from cell viability (Figure 4). We next asked whether this de-repressive effect can be related to the preferential role of Nup93-2 in NPC assembly and thus NPC levels.

To answer this question, we measured NPC levels in S2 cells depleted for Nup93-1, Nup93-2, both, Nup107 or control White, as above, using immunofluorescent staining with a classic NPC marker mAb414, combined with control Lamin Dm0 staining (Figure 5). mAb414 recognizes FG-containing Nups at several positions within the NPC and is commonly used to assess levels of mature nuclear pores [79] [80]. Depletion of Nup107, which is known to be required for NPC assembly across organisms [81], was used as a positive control. We found that similar levels of RNAi depletion as above (Figure 4) for either Nup93 paralog did not significantly alter NPC levels in S2 cells, as assessed by mAb414 staining and its fluorescent signal quantification (Figure 5A-C). In contrast, RNAi against Nup107 or both Nup93 paralogs resulted in significantly lower NPC levels, suggesting that Nup93 paralogs can compensate for each other in NPC assembly. To validate our mAb414 fluorescent signal quantification (see Methods), we also manually counted mAb414-marked nuclear foci in a single plane of nuclear surface (Figure 5C, top of cell NPC manual counts, and mean grey value assessment of the same nuclear surfaces in S5A). Our manual NPC counts similarly showed no significant difference between Nup93-1-depleted and Nup93-2-depleted S2 cells.

**Figure 5.**
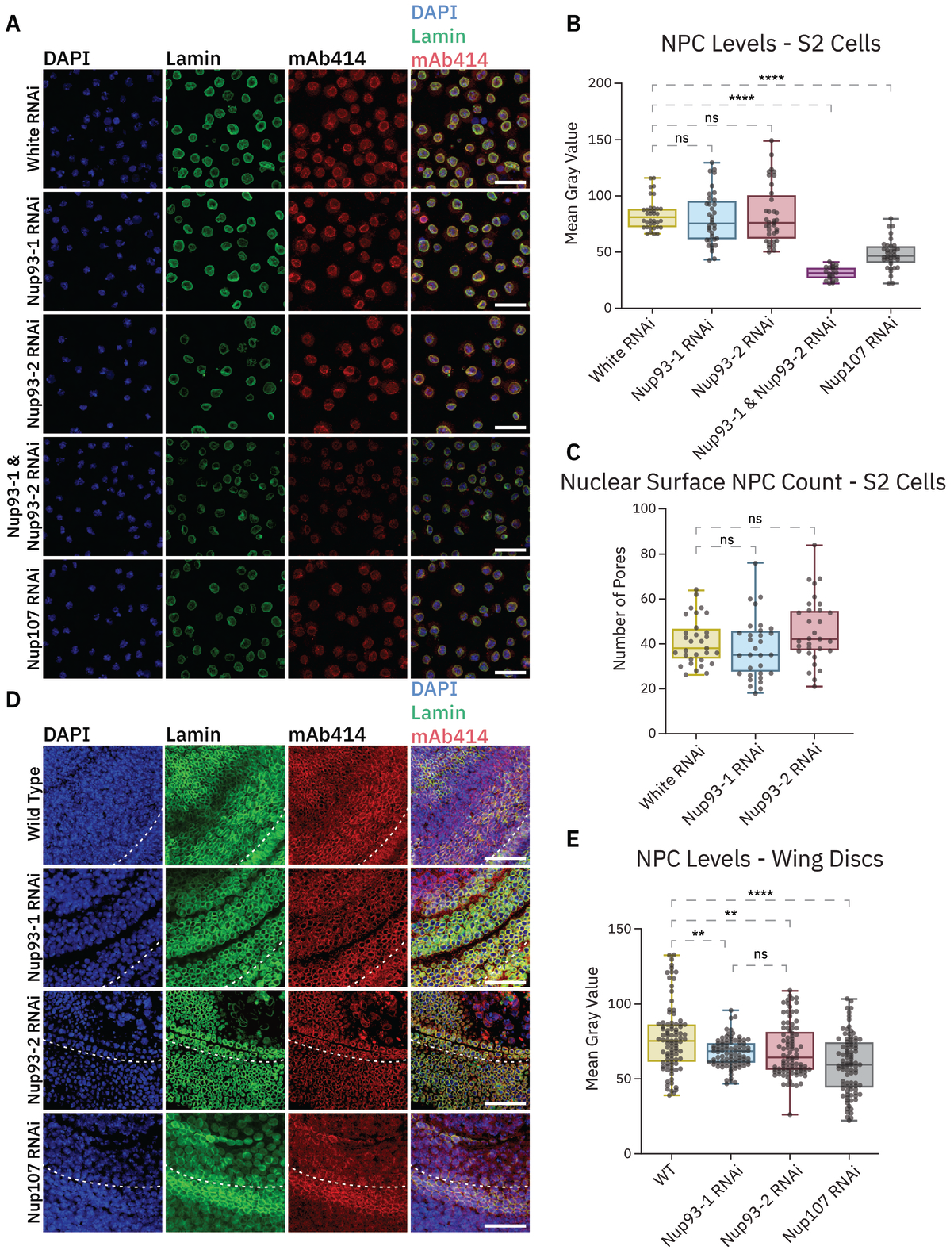
Nup93 paralogs exert similar effects on NPC biogenesis. (A) Immunofluorescent staining of S2 cells with RNAi-mediated knockdowns targeting White (negative control), Nup93-1, Nup93-2, combined Nup93-1 & Nup93-2, and Nup107(positive control). Cells were stained with DAPI (blue), Lamin (green), and mAb414 (red) to mark NPCs (63x/1.40 oil objective, 1x zoom, scale bar is 20µm). (B) Quantification of NPC levels from mAb414 signal intensity. Mean gray value of mAb414 (total pixel intensity divided by the number of pixels; n=36) was measured using ImageJ. One-way ANOVA + Dunnett’s statistical test was performed in Prism. (C) Manual quantification of NPC numbers from the top-of-cell regions (nuclear surfaces, single z slice) performed by threshold puncta counting (n=36, one-way ANOVA + Dunnett’s statistical analysis) (D) Immunofluorescent staining of larval wing discs with Nub-Gal4-driven RNAi-mediated knockdowns of Nup93-1, Nup93-2, Nup107 vs wild-type control (63x/1.40 oil objective, scale bar is 10 µm). The area above the white dashed line represents the estimated location of the knockdown in the wing blade (central area of wing pouch). Tissue was stained with DAPI (blue), Lamin (green), and mAb414 (red) for NPC levels. (E) Quantification of NPC levels from mAb414 signal intensity (mean gray value as in B) in wing discs of genotypes in (D), using ImageJ. (n=90, one-way ANOVA + Dunnett’s statistical analysis in comparison to control, unpaired t-test between Nup93-1 and Nup93-2).

We extended the analysis of NPC levels to larval wing discs, where the tumorigenic phenotype of Nup93-2 KD manifests. Unlike S2 cells, mAb414 staining of Nub-GAL4-driven Nup93-1 or Nup93-2 KD and control larval wings revealed that depletion of either Nup93 paralog led to a moderate decrease in NPC levels, yet the extent of reduction was similar between Nup93-1 and Nup93-2 KDs relative to wild-type (Figure 5D-E, S5B). Importantly, Nub-GAL4-driven Nup107 KD was found to result in a significantly more drastic reduction in NPC levels than Nup93 paralogs (Figure 5D). Since Nup107 KD does not produce a tumorigenic phenotype in the wing, these results indicate that reduction of NPC levels alone does not explain the tumorigenic phenotype of Nup93-2 KD. It also appears that Nup93 paralogs are still able to partially compensate for each other in NPC assembly in wing discs, although to a lesser extent than in S2 cells. Taken together, these results suggest that Nup93 paralogs do not exhibit drastically different roles from each other in NPC assembly and that the Nup93-2-associated tumorigenic phenotype is not explained by a unique effect of Nup93-2 on NPC levels. These findings further support the notion that de-repression of Upd genes and the associated tumor-like phenotype are based on a direct gene regulatory role of Nup93-2.

### 6. The unique tumor-like phenotype of Nup93-2 is not explained by expression differences between the paralogs

The second possibility we considered to explain the phenotypic differences between Nup93 paralogs is their differential expression in the wing. To address this possibility, we generated fly lines with individually tagged Nup93-1 and Nup93-2 using CRISPR-Cas9 (Figure 6). The Nup93-1 gene, which resides on the X chromosome, was N-terminally tagged with an HA tag, while the Nup93-2 gene, present on the 3^rd^ chromosome, was N-terminally tagged with a FLAG tag (Figure 6A-B). We confirmed the expression, correct size and specificity of the tagged Nup93 paralogs via Western blotting (Figure 6A-B). These CRISPR-tagged fly lines are homozygous viable, indicating that the inserted tags do not interfere with normal functions of these Nups.

**Figure 6.**
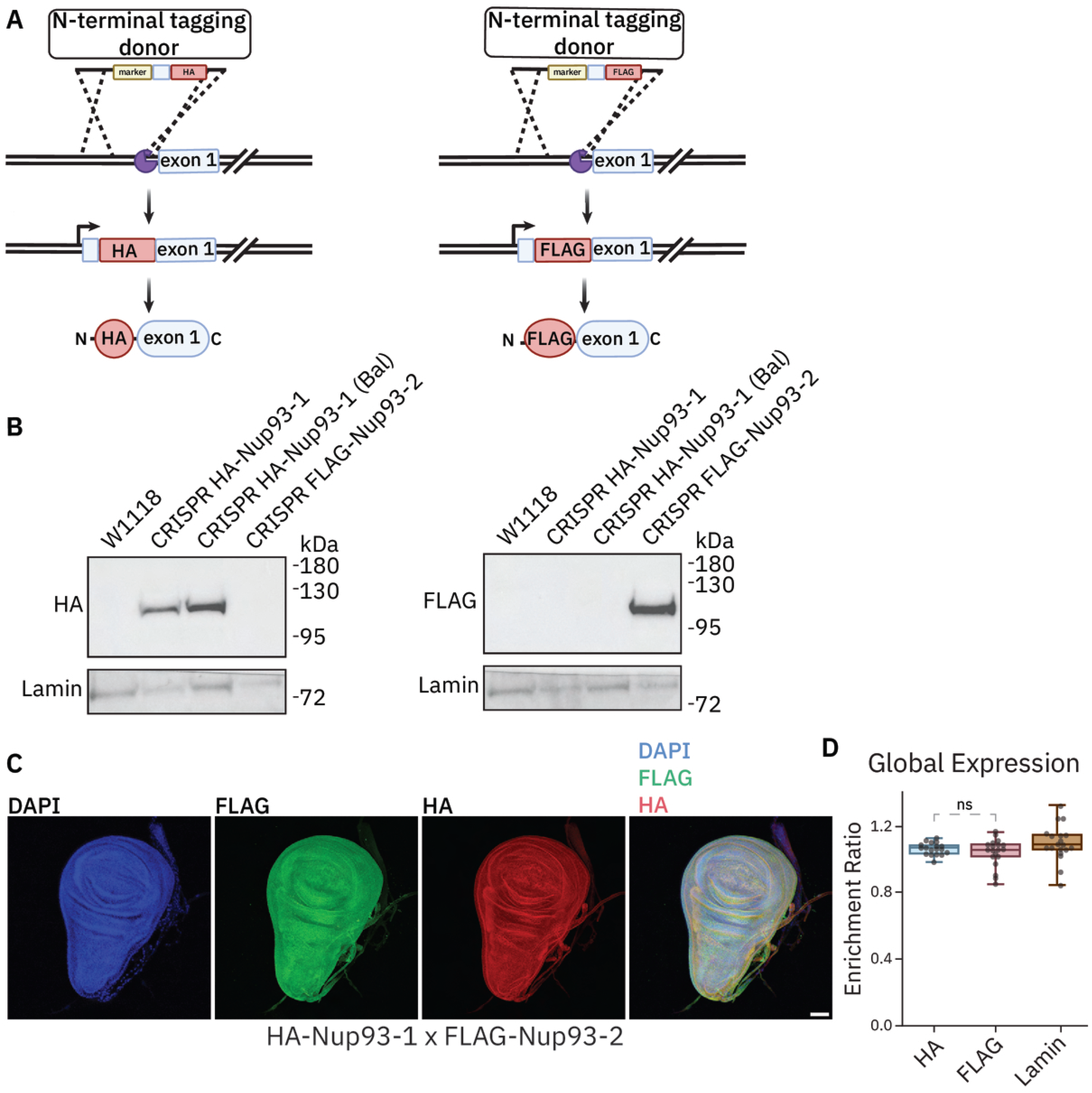
Nup93 paralogs are expressed similarly in wing imaginal discs. (A) Schematic representation of CRISPR/Cas9-mediated tagging of Nup93-1 (X chromosome) and Nup93-2 (3^rd^ chromosome) loci. Epitope tags were inserted in-frame at the N-terminus of each gene, generating HA-Nup93-1 or FLAG-Nup93-2 proteins. (made in Biorender). (B) Western blot validation of CRISPR-tagged Nup93 fly lines, using Lamin (control) and HA or FLAG antibodies. Protein lysates were obtained from female ovaries of indicated genotypes. Nup93-1 fly lines include originally obtained and balanced (Bal) lines. (C) Immunofluorescence analysis of whole-wing expression of Nup93-1 and Nup93-2, using HA and FLAG antibodies on wing discs from progeny of the HA-Nup93-1 X FLAG-Nup93-2 cross (20x/dry objective, scale bar is 50 µm). (D) Quantification of whole-wing (global) expression levels of HA-Nup93-1 and FLAG-Nup93-2 from (C), represented as a ratio of immunofluorescent signal intensity in the wing blade region to intensity in the whole wing (enrichment ratio), assessed via ImageJ. Lamin enrichment is also shown for comparison. Note no significant difference between Nup93-1 and Nup93-2 (n=10).

To assess tissue-specific expression, we assessed endogenous expression of Nup93-1 and Nup93-2 using HA and FLAG immunofluorescent staining (Figure 6C-D, S6A-B). As we were primarily interested in wing phenotypes, we analyzed HA and FLAG distribution and levels in wing discs from animals carrying both tags (which resulted from crossing the two lines to each other). We detected expression of both Nup93 paralogs throughout the wing disc and did not observe any notable differences in wing expression patterns (Figure 6C-D). Importantly, we did not detect any preferential enrichment of either tag in the blade/pouch region of the wing, where the tumorigenic phenotype of Nup93-2 manifests (Figure 6D). Similarly, analysis of HA and FLAG tagged Nup93 paralogs in adult ovaries and in larval salivary glands did not show any obvious differences in global expression levels (Figure S6A-B). We also observed localization of both HA and FLAG to nuclear rims, as marked by nuclear pore markers mAb414 or an antibody to another core NPC subunit Nup154, in all analyzed tissues (Figure S6A-C, S6C shows validation of generated antibody to Drosophila Nup154), demonstrating the expected cellular targeting of Nup93 paralogs. Although additional expression analysis may shed light on other tissue-specific requirements of Nup93 paralogs (Figure 1B), we concluded that the Nup93-2-specific wing phenotype is not explained by differential expression patterns of Nup93 paralogs in the wing.

### 7. Nup93-2 exhibits a unique NE localization pattern in a tissue-specific manner

A third possibility we considered to explain the phenotypic differences of Nup93 paralogs is the potentially unique sub-cellular localizations of Nup93-2 and Nup93-1 relative to the NPC. As both Nup93 paralogs are clearly identifiable as homologs of mammalian Nup93 and yeast Nic96, they are predicted to occupy similar positions within the NPC structure as components of the Nup93-Nup154 inner-ring sub-complex. We wondered however if Nup93-1 and Nup93-2 1) may localize to NPCs in a mutually exclusive way or co-localize at the same NPCs, and 2) are targeted exclusively to NPCs or localize to any additional sub-nuclear structures. To investigate these questions, we initially co-transfected S2 cells with generated constructs of Nup93-1-Myc and Nup93-2-GFP and analyzed their potential co-localization with each other and with mAb414-marked NPCs (Figure S7A-B). We found that Nup93-1 and Nup93-2 exhibited high levels of co-localization with each other and with mAb414 in S2 cells, with Pearson Correlation Coefficient (PCC) of 0.8 and above. We did notice a slightly lower co-localization of Nup93-2 with NPCs, but overall, these results suggested that Nup93 paralogs can be assembled into the same nuclear pores, which is not unexpected.

To analyze localization of endogenous Nup93 paralogs, we utilized our CRISPR-tagged fly lines of HA-Nup93-1 and FLAG-Nup93-2 (Figure 6). Similarly to S2 cells, we compared localization of each Nup93 paralog to a component of the NPC and to each other, using HA and FLAG co-staining with mAb414 and Nup154, in larval wing discs (Figures 7A-B). Surprisingly, we observed that although HA-Nup93-1 co-localized nearly perfectly with mAb414-marked mature NPCs, FLAG-Nup93-2 showed a strikingly partial co-localization with Nup93-1-HA (Figure 7A). To quantify the extent of co-localization, we utilized the method our lab developed previously [82,83] to determine PCC values of pixel-by-pixel correlation between 2 fluorescent signals. Using this method, we obtained expectedly high correlation values of ∼0.8 for HA-Nup93-1 and mAb414 co-localization, but only moderate values of ∼0.55 for FLAG-Nup93-2 and HA-Nup93-1 co-localization (Figure 7B), suggesting that Nup93-2 may target an additional location within the nuclear periphery in cells of the wing disc. On the other hand, FLAG-Nup93-2 showed much higher co-localization with Nup154, with PCC values of ∼0.7, yet Nup154 co-localized at similarly low levels with mAb414, with PCC values of ∼0.45 (Figure 7A-B). Based on this analysis, we conclude that unlike Nup93-1, Nup93-2 together with Nup154 may form a separate sub-nuclear complex in addition to normal NPCs.

**Figure 7.**
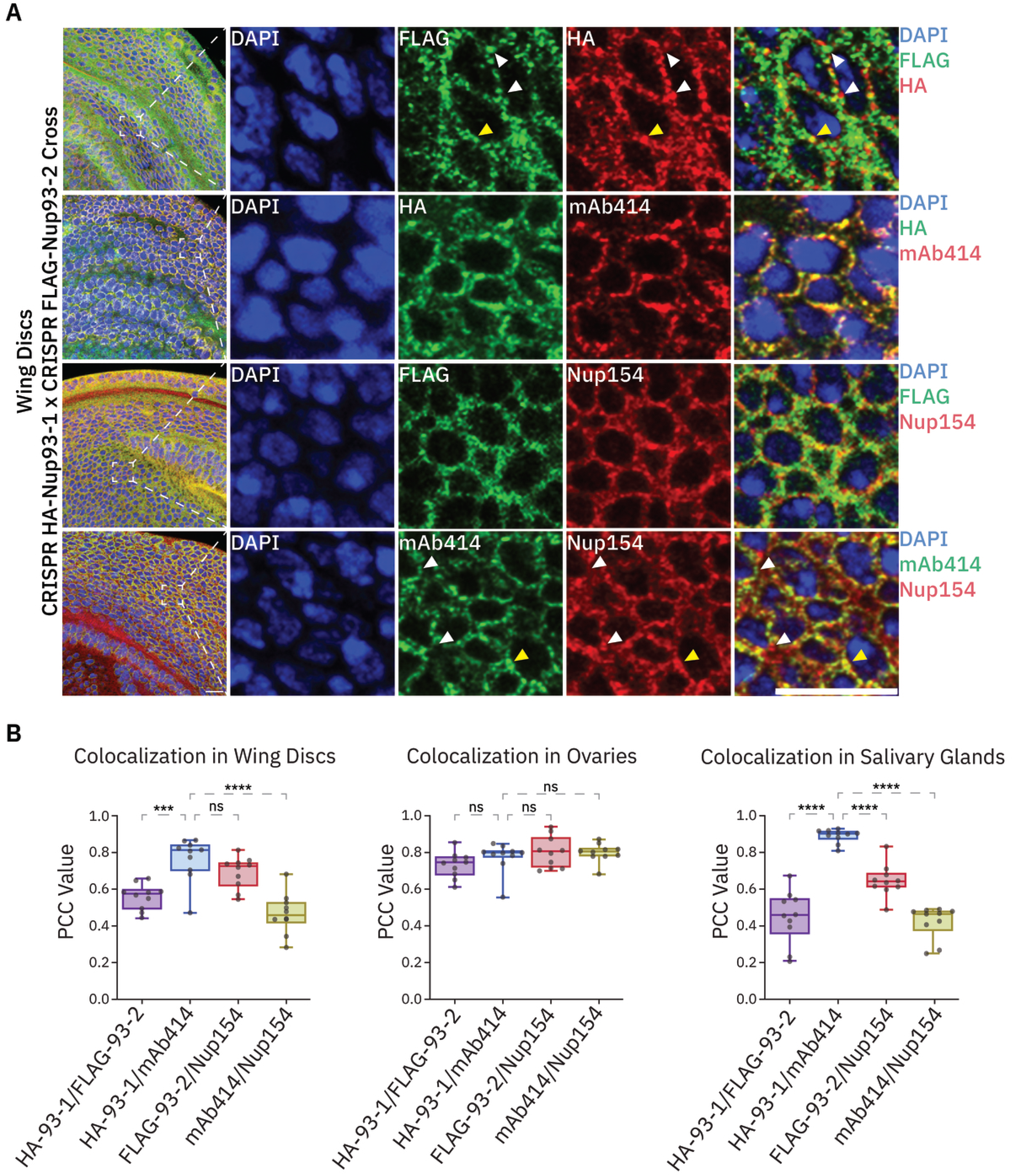
Nup93-2 localizes to non-NPC locations in a tissue-specific manner. (A) Immunofluorescence analysis of Nup93-1 vs Nup93-2 localization, using HA, FLAG, mAb414, and Nup154 antibodies in wing discs of larvae carrying both HA-Nup93-1 and FLAG-Nup93-2, as indicated. Left column shows entire obtained images, with outlined locations of zoomed-in images that are shown to the right. White arrows point to examples of unique signal, while yellow arrows point to examples of overlapped signal (63x/1.40 oil objective, scale bar is 10 µm). Note nearly perfect co-localization between Nup93-1 and mAb414. (B) Analysis of co-localization of each Nup93 paralog, using HA or FLAG antibodies, with NPC marker mAb414 or antibody to Nup154 in indicated tissues, using the PCC (Pearson Correlation Coefficient) pixel-by-pixel method, performed in MATLAB (see Methods). N=10 for each tissue, one-way ANOVA + Dunnett’s statistical analysis in Prism.

Interestingly, an NPC-related but distinct nuclear complex containing the Nup154 yeast homolog Nup170 has been described [84]. In budding yeast, Nup170 binds sub-telomeric heterochromatin and interacts with Sir4, a component of the silent information regulator (SIR) heterochromatin-binding complex [34]. Nup170 and Sir4 were found to interact with additional components and importantly, a subset of core Nups, forming what was termed the Sir4-associated Nup (Snup) complex [84]. The Snup complex, located at the NE but shown to be distinct from classic mature NPCs, was found to contain the Nup93 yeast homolog Nic96 but not many of the auxiliary FG-containing Nups, which parallels the observed lower co-localization between Drosophila Nup154 and by extension, Nup93-2 with mAb414 in our system. Our co-localization analysis suggests that in the wing disc, Nup93-2 may be part of both NPCs and the Drosophila equivalent of the Snup complex, while Nup93-1 is found exclusively at NPCs.

To understand the generality of this phenomenon, we extended our co-localization analysis to two other tissues – the developing egg chambers within female ovaries and the salivary glands of third instar larvae, which unlike wing discs, contain post-mitotic polytenized cells. Surprisingly, we found that the correlation values between all tested factors were high in the ovaries, indicating a high level of co-localization between Nup93-1 and Nup93-2, Nup154 and mAb414, Nup93-1 and mAb414, and Nup154 and Nup93-2 (Figure 7B). On the other hand, nuclei of the larval salivary glands exhibited a very similar pattern of co-localization to larval wing discs, such that Nup93-2 and Nup154 once again showed only partial co-localization with Nup93-1 and mAb414 (Figure 7B). These results suggest that the putative Snup complex forms in the salivary gland but not in the ovary, indicating tissue-specific regulation of Nup93-2 localization. Together, our findings identify a distinct sub-nuclear distribution of Nup93-2 relative to Nup93-1 and suggest an intriguing possibility that the unique gene silencing function of Nup93-2 may involve a separate Nup-containing complex that targets a subset of Polycomb-repressed genes, some of which influence tissue growth (Figure 8, model).

**Figure 8.**
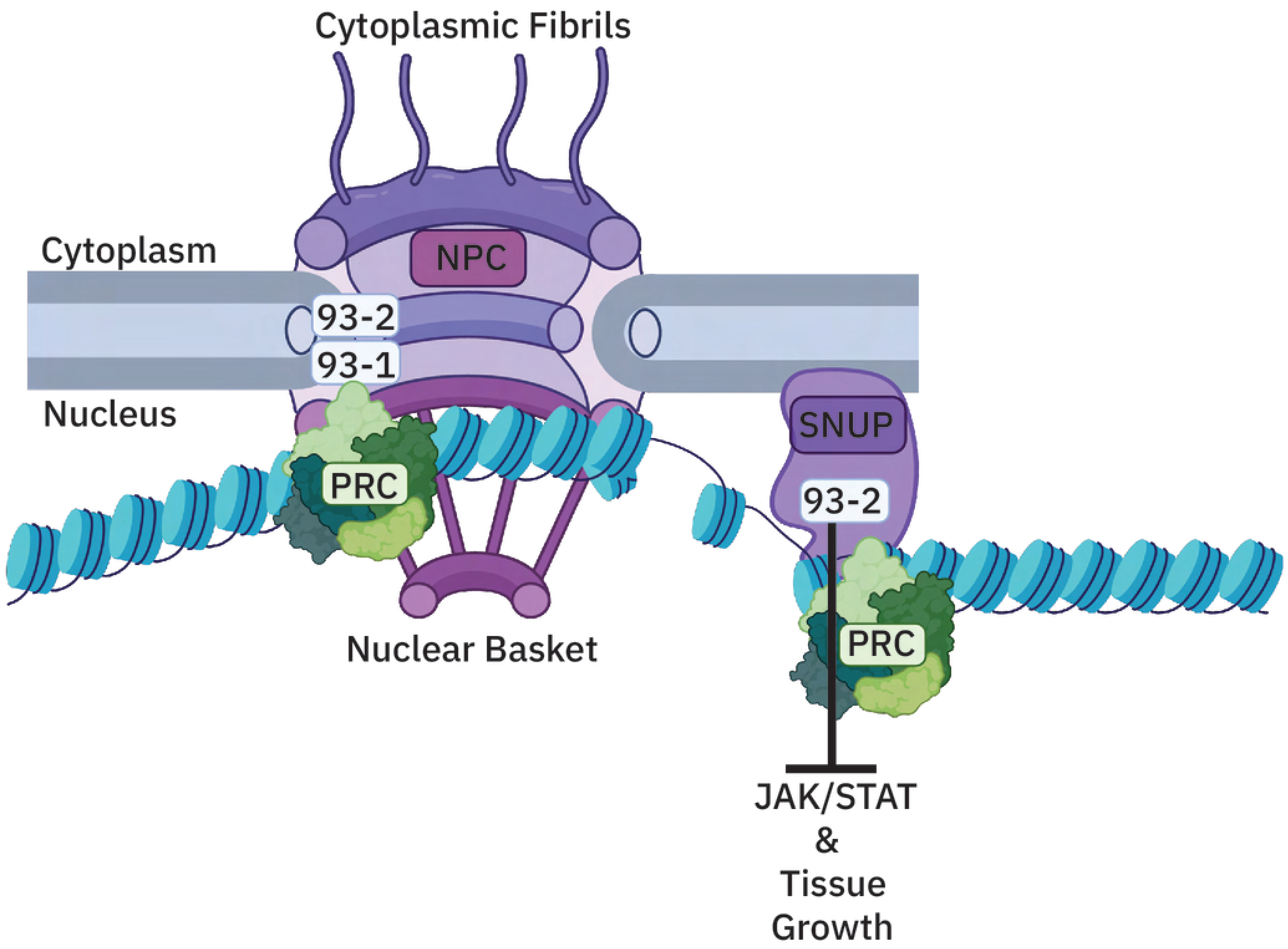
Nup93-2 paralog facilitates Polycomb-mediated gene silencing of growth-promoting genes - a putative model. A schematic of a working model on the roles of Nup93 paralogs in gene silencing and growth regulation. While both paralogs can likely associate with Polycomb domains at the canonical NPCs, Nup93-2 may be present at non-NPC locations as part of Snup-like complexes that interact with a subset of Polycomb domains that normally repress JAK/STAT ligands and other growth-promoting genes. Knockdown of specifically Nup93-2 in wing discs compromises Polycomb-mediated silencing of the JAK/STAT pathway, leading to tumor-like tissue overgrowth. PRC stands for Polycomb Repressive Complex (green). String of blue spheres represent chromatin. The Snup complex is putatively portrayed as associated with the inner nuclear membrane of the NE.

## DISCUSSION

NPC components are known to regulate gene expression and tissue-specific development, via both their transport-dependent and chromatin-binding functions [85,86]. We and others have previously identified binding of NPC components to facultative heterochromatin [37,39], but the consequences of that relationship during in vivo development have been poorly defined. Here, we investigated tissue-specific roles of two distinct Drosophila Nup93 paralogs and their unique functions in regulation of gene expression and NPC assembly. We found that different tissues exhibit unique requirements for Nup93 paralogs and that unlike Nup93-1 and other tested Nups, Nup93-2 depletion in wing imaginal discs leads to a tumor-like overgrowth phenotype. Our RNA-seq analysis demonstrated a widespread gene upregulation in Nup93-2-depleted tumors and revealed a particularly drastic de-repression of the Upd gene cluster as well as associated hyper-activation of JAK/STAT signaling. This phenotype is reminiscent of imaginal disc tumors produced by Polycomb mutations and which are similarly driven by upregulation of JAK/STAT signaling [56,64,66]. Consistently, we found that Upd and Wg/Wnt gene clusters reside within Polycomb domains and correspond to binding peaks of pan-Nup93 ChIP-seq, produced with an antibody that likely recognizes both Nup93 paralogs. Importantly, RNAi depletion of Nup93-2 in S2 cells also resulted in strong de-repression of Upd genes, indicating that this de-repression is likely not an indirect consequence of tissue damage.

Overall, these results particularly implicate the Nup93-2 paralog in Polycomb-mediated silencing and suggest that the role of Nup93-2 in JAK/STAT upregulation and tumor growth stem from direct binding to Polycomb domains.

We found that a differential effect on NPC biogenesis does not explain the phenotypic differences between the Nup93 paralogs or the tumorigenic phenotype of Nup93-2. No significant differences in NPC levels were observed between knockdowns of the two paralogs in either wing discs or S2 cells. Furthermore, RNAi knockdown of Nup107, a structurally indispensable Nup, lead to more significantly reduced NPC levels yet did not cause a tumorigenic phenotype. We also did not observe significant differences between the two paralogs on cell viability in S2 cells, further indicating that their effects on NPC assembly are comparable. This likely suggests that Nup93-1 and Nup93-2 can fully or partially compensate for each other as structural components of the NPC. Recent structural studies in vertebrate cells revealed several locations of Nup93 in addition to the classically known inner-ring position [87], and whether Drosophila Nup93 paralogs can differ in their preferred locations within the NPC remains to be determined. Moreover, analysis of Nup93-1 and Nup93-2 expression levels using CRISPR-inserted tags did not reveal notable differences in the wing disc. The full spectrum of their expression differences across tissues awaits future analysis, but in the present study, paralog-specific expression differences in the wing do not explain the unique phenotype of Nup93-2 KD.

The main difference that we identified between Nup93 paralogs is their sub-nuclear localization. While we observed expectedly high levels of co-localization between Nup93-1 and mAb414, our analysis identified a unique nuclear pool of Nup93-2 that does not co-localize with Nup93-1 and by extension, with mAb414-marked mature NPCs. Interestingly, this pool of Nup93-2 co-localizes to a higher extent with NE-associated Nup154, another inner-ring sub-complex component. Consistently, Nup154 also shows only partial co-localization with mAb414-marked NPCs, and the extent of this co-localization parallels co-localization levels of Nup93-2 with Nup93-1. These results support an intriguing notion that Nup93-2 and Nup154 may form a unique NE-associated complex, separate from the canonical NPCs. This complex is highly reminiscent of an NPC-independent Snup complex identified in yeast, which contains homologs of both Nup154 and Nup93, as well as other core Nups, yet lacks many FG Nups [84].

Interestingly, Drosophila Nup154 has been suggested to mediate chromatin attachment in a way that can be suppressed by Nup93-1 and FG-containing Nup62 [88], suggesting that its chromatin-binding activity can be regulated by specific Nup partners. Furthermore, the NPC-targeting domain of Nup154 was shown to not be required for its identified role in stabilization of select INM proteins at the NE [89], which may indicate two separate functions of Nup154 at the NPC vs the NE. Both of these studies support the proposed existence of a Snup-like chromatin-binding complex in Drosophila cells. As opposed to Sir-4-bound sub-telomeric heterochromatin in yeast, our results suggest that such a complex in Drosophila may have evolved interactions with Polycomb-bound heterochromatin.

It is also interesting that potential existence of a Snup-like complex, as revealed by only partial co-localization of Nup93-2 and Nup154 with NPCs, appears to be tissue-specific. Whereas in S2 cells and oocytes, Nup93-2 and Nup93-1 were found to co-localize with each other and mature NPCs at a high level, in larval tissues such as imaginal discs and salivary glands this was not the case. This variability suggests that in Drosophila, formation of an additional Nup93-2/Nup154 Snup-like complex may be developmentally regulated – absent in early stages such as oocytes and embryonic cells, but forms more readily in more differentiated cell types. Alternatively, the Snup-like complex may form exclusively in specific cell types, independently of the organismal developmental stage. Because both salivary gland and late-stage nurse cell nuclei of egg chambers are non-dividing, cell cycle stage is unlikely to account for the difference in co-localization. It is also worth noting that high levels of co-localization in developing egg chambers suggest that our observed co-localization differences do not stem from technical limitations.

Together, our combined results lend to a model where Nup93-2-containing Snup-like complex at the nuclear periphery binds a subset of Polycomb domains that silence growth-promoting genes such as the Upd gene cluster (Figure 8). At such genes, Nup93-2 facilitates Polycomb-mediated silencing during wing development, such that loss of Nup93-2 leads to their significant upregulation and resulting tumor-like overgrowth. However, many questions remain. Although our RNA-seq and RT-qPCR analysis identified a stronger silencing function for Nup93-2, both Nup93 paralogs and both canonical NPCs and a Snup-like complex may target Polycomb domains (Figure 8). Knockdown of Nup93-1 exhibits an overlapping set of de-repressed genes with knockdown of Nup93-2, suggesting that Nup93 paralogs have both shared and unique targets. Interestingly, unique DEGs of Nup93-1 showed enrichment for genes involved in neuronal differentiation, pointing to a potential tissue-specific gene regulatory role of Nup93-1. This role is supported by the unique requirement of Nup93-1 in developing neurons (Figure 1). Whether these genes represent direct binding targets of specifically Nup93-1 remains to be determined. Similarly, mapping chromatin binding of specifically Nup93-2 will be highly informative and allow us to further investigate its unique binding targets. Based on existing data, we hypothesize that Nup93-1 and Nup93-2 bind both shared and unique target genes and Polycomb domains, with Nup93-2 playing a critical role in silencing of genes required for growth control.

It is also likely that a proportion of the differentially expressed genes we observed in Nup93-1 and Nup93-2 KDs arise from activating roles of NPC components or from their indirect roles related to nuclear transport. In both flies and mammalian cells, Nups have also been found to associate with enhancers and super-enhancers [90]. Acute depletion of Nup93 in mammalian cells led to both downregulation and upregulation of Nup93-targeted genes [91], revealing the ability of Nup93 to facilitate both activities. It is tempting to speculate that in Drosophila, Nup93-2 plays a preferential gene silencing role at Polycomb domains, some of which are specifically targeted to a potential Snup-like complex, while Nup93-1 can toggle between activating and silencing roles, depending on context, at canonical NPCs. In this manner, Nup93-2 may function as a consistent silencing module, reserved for a subset of Polycomb domains that require terminal or robust silencing. Nup93-1 on the other hand, may also be connected to reported roles of Polycomb machinery at active genes, where PcG proteins appear to have architectural functions in scaffolding chromatin loops [92]. Architectural functions have also been linked to multiple NPC components [28,35,37] [93], including at Polycomb targets, and any distinct roles of Drosophila Nup93 paralogs in genome architecture await further investigation.

Our findings can further inform human pathologies that are linked to Nup93 and Nup155, and to NPCs in general. Human Nup93 has been repeatedly implicated in oncogenesis, with aberrantly high levels of Nup93 implicated in cancer progression and aggressiveness [94]. For example, elevated levels of human Nup93 in bladder cancer correlated with poor prognosis, and knocking down Nup93 in bladder cancer cell lines decreased cell proliferation and downregulated Wnt signaling [53]. In our study, we identified an opposite effect of Nup93-2, knockdown of which resulted in tissue overgrowth and increased Wnt/Wg signaling. This suggests that Drosophila Nup93-2 may have evolved an inverse function in growth control, which points to an overall relationship between Nup93 homologs and growth-promoting pathways. Interestingly, another Nup-associated wing disc tumor phenotype, produced by Nup98-Nup96 depletion combined with an apoptosis inhibitor, was suggested to arise primarily via defective protein synthesis due to compromised nuclear export of ribosomal protein Rpl10Ab [77]. Although it remains to be determined whether Nup93-2 loss-induced tumors similarly involve defects in protein synthesis, our results suggest that Nup93-2-associated tumors may arise via a distinct chromatin-based mechanism. It is also of note that Nup93-2-associated tumors arise solely from the loss of Nup93-2, with no additional requirement for inhibition of apoptosis. Conversely, such tumors may be based on multiple mechanisms happening in parallel, with compromised Polycomb-mediated gene silencing as one key driver. Together, these results highlight the importance of NPC components in regulation of cell proliferation, signaling pathways and gene expression that likely contribute to a wide range of physiological processes.

## MATERIALS AND METHODS

### Fly maintenance, genetics and stocks

All stocks and crosses were raised at RT or 25C on standard fly food (Fisherbrand Jazz-mix Drosophila food), and crosses were flipped every 4-5 days. The following genetic stocks were used – BDSC 34090 (Nup93-1 RNAi), BDSC 51758 (Nup93-2 RNAi), VDRC 22407 (Nup107 RNAi), BDSC 26197 (10XStat92E-GFP), BDSC 86108 (Nub-Gal4, wing discs and salivary glands), BDSC 5534 (ey-Gal4, eye discs), BDSC 7011 (cg-Gal4, fat body), BDSC 35695 (Nup62 RNAi), BDSC 58290 (Nup133 RNAi), BDSC 34580 (Nup50 RNAi), BDSC 55375 (Nup42 RNAi), BDSC 65219 (Nup210 RNAi), VDRC 24265 (Mtor RNAi). The Mef2-Gal4 (muscle) stock was gifted by lab of Dr. Richard Cripps, Elav-Gal4 (neurons) and Repo-Gal4 (glia) lines were gifted by Dr. Kim Finley.

CRISPR-Cas9 insertions of FLAG and HA were designed to the N-termini of Nup93-2 and Nup93-1, respectively, and CRISPR fly lines were generated via WellGenetics Inc. CRISPR-tagged fly lines were crossed to each other so both HA and FLAG tags were present in the F1 progeny, which was used for analysis. To assess STAT-GFP, a fly line carrying both 10XStat92E-GFP and UAS-Nup93-2 RNAi was generated via genetic crossing, then crossed to Nub-Gal4. As a control, the 10XStat92E-GFP line was crossed to Nub-Gal4.

### RNA-Seq

To generate wing-specific knockdowns, Nub-Gal4 were crossed to UAS-RNAi lines of Nup93-1, Nup93-2 or to control w1118. Large-scale crosses of 15-25 females with a ratio of 2.5:1 females to males were raised at 25°C, set up in triplicate per genotype. Crosses were flipped on day 4, and wing imaginal discs from L3 larvae were harvested on day 6. Dissections per genotype were performed within 1 hour and were immediately placed in a 1.5mL centrifuge tube and frozen using liquid nitrogen, then placed at −80°C. RNA was extracted from collections of 40 wing discs per genotype per replicate, using Qiagen RNAeasy Mini Kit. Prepared RNA samples were sent to Novogene, Inc. for QC, library prep, and sequencing.

### RNA-seq bioinformatics analysis

Paired-end RNA-sequencing reads were assessed for quality with FastQC (v0.11.8) and adapter- and quality-trimmed with Trim Galore (v0.6.6) using Phred quality cutoff 20. Trimmed reads were aligned to the *Drosophila melanogaster* genome (BDGP6.32/dm6, Ensembl release 107) with STAR (v2.7.5b) in two-pass mode: a first pass per sample collected novel splice junctions, which were merged across all samples and supplied to a second pass (-- outFilterMultimapNmax 20, non-canonical junctions removed). Alignment quality was evaluated with samtools (v1.11), Qualimap (v2.2.1) and Preseq (v2.0), and all preprocessing metrics were aggregated with MultiQC (v1.8). Gene level read counts were obtained by using Salmon (v1.2.1, RRID:SCR_017036) for all libraries [95]. Sample normalization was carried out using the median-ratios normalization method from DESeq2 R package (v1.30.1, RRID:SCR_015687) [96]. Gene-level counts were rounded to integers, and genes with fewer than five reads summed across samples were removed (12,340 genes retained). Differential expression was performed with DESeq2 (v1.40.2; R v4.3.1): the three wing genotypes (driver-only control, Nup93-1 RNAi, Nup93-2 RNAi; n = 3 each). Genes with a Benjamini–Hochberg-adjusted *p* < 0.05 and |log2 fold change| > 1 were considered differentially expressed. Volcano plots (Glimma v2.10.0) display apeglm-shrunken log2 fold changes on the x-axis against the −log10 *p*-value on the y-axis, with points colored by differential-expression status (adjusted *p* < 0.05 and |log2 fold change| > 1) (Fig. 2A,B). For the union of differentially expressed genes across both knockdowns, variance-stabilizing-transformed (VST) counts were row-wise z-scored and displayed as a heatmap (heatmaply v1.4.2) with genes clustered by Euclidean distance and complete linkage and samples held in genotype order (Fig. 2C). Knockdown specificity was confirmed by comparing VST-normalized expression of each *Nup93* paralog across the three genotypes (Fig. 2D). Overlap of up- and down-regulated genes between the two knockdowns was visualized as proportional Venn diagrams (Fig. 2E), and the corresponding gene lists were submitted to FlyEnrichr (https://maayanlab.cloud/FlyEnrichr) for functional enrichment (Fig. 2E). Fig. 2E lists top enriched categories (by combined score) from pathways or function analysis via WikiPathways2018, Anatomy AutoRIF, and Allele LoF Phenotypes from FlyBase 2017. Lists of DEGs for each condition can be found in Tables S2 and S3. For comparisons to ChIP-seq datasets at Upd and Wg loci, the following public datasets from published work have been used: ChIP-seq datasets of Pc and H3K27Me3 (wing discs, GSM3424769 and GSM3424770, from GSE121028) and of pan-Nup93 (S2 cells, GSE135610).

### S2 cells culture and transfections

Drosophila S2 cells were grown at 25°C in Gibco Schneider’s *Drosophila* media (ThermoFisher Scientific) with 10% FBS and 1X Penicillin/Streptomycin. For transfections, cells were plated on a 6 well dish, allowed to adhere overnight and grown to approximately 1 x 10^6^cells/mL. Media was changed to 1mL of serum-free, and a transfection mixture containing 100µL of serum-free media and a 3:2 ratio of DNA to Fugene (commonly, 3μg of DNA and 2μL of Fugene HD) was added to the wells. After incubation at RT for 3 hours, 2mL of serum-containing media was added to each well. Transfected cells were incubated at 25°C for 72 hours, then harvested and assessed by IF or WB.

### S2 cells RNAi knockdowns

Double-stranded RNA (dsRNA) was prepared for white, Nup93-1, Nup93-2, Nup107, and ph-p using Invitrogen MEGAscript T7 *in vitro* transcription kit (ThermoFisher Scientific), using primers listed in Table S1, according to manufacturer’s instructions and similarly to [37]. S2 cells were plated at ∼1.5 × 10⁶ cells per well in a 6-well plate and washed in PBS and resuspended in serum-free media before dsRNA treatment. dsRNA targeting the gene of interest was added at 12 µg per 800 µL per well. Cells were incubated with dsRNA for 1-2 hours at 25°C before adding complete media. They were then maintained at 25°C for 72 hours. A second or third dsRNA treatment were performed under similar conditions on days 3 and 5, with adjustment of cell density to the starting numbers. Cells were collected on days 1-7, with day 7 being the most common collection point for most experiments. Knockdown efficiency was validated by RT-qPCR.

### Antibodies and DNA constructs

The following primary antibodies were used – mAb414 (Covance mouse monoclonal MMS-120P), anti-Lamin Dm0 (rabbit polyclonal antiserum #837 gifted by Dr. Paul Fisher), anti-HA (Santa Cruz mouse monoclonal F-7 and DSHB rabbit anti-HA rRb-IgG), anti-FLAG (Sigma Aldrich mouse monoclonal M2 antibody F1804), anti-GFP (Abcam rabbit polyclonal ab290), anti-beta tubulin (DSHB mouse monoclonal E7), anti-Wg (DSHB Wingless-AB_528512), anti-Myc (Santa Cruz mouse monoclonal 9E10). Secondary antibodies included Abcam goat anti-rabbit ab205718, Abcam goat anti-mouse ab205719, Abcam goat anti-mouse Alexa Fluor® 488, Abcam goat anti-mouse Alexa Fluor™ 546, ThermoFisher goat anti-rabbit Alexa Fluor™ 488, Abcam goat anti-rabbit Alexa Fluor® 568.

To generate anti-Nup154 antibody, a fragment of Nup154 cDNA corresponding to amino acids 561-770 was cloned into pETG-41A vector via Gateway cloning. The construct was transformed into BL21 bacterial cells, and expression was induced with 0.1mM IPTG. The resulting recombinant His-MBP-Nup154 protein was purified with a HIS tag using Ni-NTA agarose beads and sent to Pacific Immunology Corp. for rabbit antibody generation and subsequent affinity purification. Generated antisera were validated by RNAi and western blotting in S2 cells.

The following cDNA constructs were obtained from DGRC and used as templates for cloning or dsRNA generation – LD21129 (Nup93-1), GH01807 (Nup93-2), LD18761 (Nup107), LD01444 (ph-p), GH19653 (White), LD21772 (Nup154). Nup93-1 and Nup93-2 cDNAs were inserted into pAFMW and pAGW vectors, respectively, via Gateway cloning for transfection into S2 cells.

### S2 Cells Immunofluorescence

S2 cells (treated with RNAi or transfection) were collected, pelleted, and resuspended in PBS before being plated onto poly-L-lysine–coated slides and allowed to adhere for 30 minutes. Slides with adhered cells were then rinsed in PBS, and cells were fixed 2% paraformaldehyde in 0.1% PBS+0.1%TritonX (PBSTx) for 7-10 minutes. After fixation, cells were further permeabilized and washed in PBSTx 3 times for 5 minutes each, then blocked in PBSTx+1% BSA (blocking solution) for 30 minutes. Primary antibodies incubation was performed in blocking solution in a humidified chamber at 4°C overnight. Next day, slides were washed in PBSTx as before and incubated with secondary antibodies (diluted 1:500) in the humidified chamber for 2 hours at RT in the dark, then washed 3 times in PBSTx, stained with DAPI (diluted 1:1000 in PBS), washed again in PBS and allowed to completely dry. Vectashield was added before adding the coverslis, after which slides were sealed and used for confocal microscopy imaging.

### Image analysis – S2 cells

All slides were imaged on a Leica STELLARIS confocal microscope at the same zoom (1x) and consistent intensity for each channel (DAPI, Lamin, mAb414). Images were imported into ImageJ, and the red channel (mAb414) was used for analysis. To quantify NPC levels via mAb414 staining, images were max-projected from entire Z stacks and analyzed for mean gray value (total pixel intensity divided by area). Triplicate RNAi experiments with two technical replicates per each RNAi treatment were used, and 6 cells were selected per slide, yielding 36 cells per treatment. Obtained mean gray values were graphed and analyzed in Prism, with a one-way ANOVA + Dunnett’s statistical test. An alternative manual quantification was done to validate results by only using a single top z-stack slice for each image and counting manually by hand for puncta (representing nuclear pores). For each condition, triplicate RNAi experiments with two technical replicates per each RNAi treatment were used, and 6 cells were selected per slide (as above). Obtained manual counts were similarly analyzed in Prism. To assess co-localization of Nup93 paralogs with each other and with mAb414 in transfected cells, slides were analyzed in ImageJ with the BIOP JACoP plugin. Images were split into red and green channels, and a manual threshold was applied. Output of the Pearson Correlation Coefficient was recorded, and graphs were created for these values.

### Fly tissues Immunofluorescence

Wandering L3 larvae were dissected for wings and salivary glands, and adult female flies were dissected for ovaries in PBS. Tissues were fixed in a PBS+4% PFA solution for 10 minutes. Tissues were then rinsed in PBS for 10 minutes, washed 3 times for 5 minutes each in a PBS+0.3% TritonX (0.3% PBSTx) solution. 300 μL of primary antibodies diluted in 0.3% PBSTx were added to the tissues, which were then incubated at 4°C for up to 3 days. Tissues were washed in 0.3% PBSTx as before. Secondary antibodies (diluted 1:500 in 0.3% PBSTx + 1% BSA) were added, and tissues were incubated on a shaker for 2-4 hours or overnight in the dark at RT. Tissues were then washed 3 times for 5 minutes each in 0.3% PBSTx. DAPI was applied at 1:1000 dilution, and tissues were rinsed in 0.3% PBSTx, mounted on slides with 15μL of Vectashield, covered with coverslips, and imaged using confocal microscopy.

### Image Analysis – fly tissues

All slides were imaged on a Leica STELLARIS confocal microscope at the same zoom (1x) and consistent intensity per each channel. Images were analyzed in ImageJ. The images of wing disc pouch/blade were sub-divided into the wing blade edge, the central wing blade, and the wing blade/hinge regions, to ensure accurate quantification across the RNAi-affected region. 10 cells per region were quantified, for a total of 30 cells per wing. The cells were manually traced for the outer ring and the inner ring of nuclei, using Lamin and DAPI as markers, and measured values were exported into Excel to calculate the mean gray value of the nuclear periphery. The final mean gray value was then determined by subtracting the integrated density of the inner from the outer ring values and dividing by the corresponding area. These values were plotted and analyzed in Prism - two statistical tests were applied, including one-way ANOVA + Dunnett’s statistical analysis in comparison of all to control genotype and an unpaired t-test for differences between Nup93-1 and Nup93-2.

For whole-wing (global) expression analysis, wing discs of F1 progeny from the CRISPR-tagged lines cross were stained for FLAG and Lamin, HA and Lamin, and HA and FLAG, and the resulting images were analyzed in ImageJ. For FLAG, HA, or Lamin, an enrichment ratio was calculated using the mean gray value of the wing blade (pouch) divided by the mean gray value of the whole wing. These values were analyzed in Excel and Prism using an unpaired t-test. To quantify STAT-GFP expression levels, 10XStat92E-GFP; Nup93-2 RNAi fly line was crossed with a Nub-Gal4 driver, and wing discs of F1 progeny, stained for GFP and Lamin, were imaged and analyzed in ImageJ. Mean gray values of GFP signal were quantified in the inner and outer wing blade (pouch). The inner value was divided by the outer value to produce a STAT-GFP enrichment ratio (which is normally enriched in the outer part of the wing pouch). The values were analyzed in Excel and Prism using an unpaired t-test to compare between 10XStat92E-GFP alone control and 10XStat92E-GFP; RNAi Nup93-2.

### MATLAB Colocalization Quantification

The MATLAB code used for colocalization analysis in fly tissues was adapted from the one previously described [82,83]. Colocalization analysis between HA-Nup93-1, FLAG-Nup93-2, mAb414, and Nup154 was performed in ovaries, wing discs, and salivary glands. Images for each combination per tissue were acquired for each channel along with a nuclear stain (DAPI). Channels were separated and saved as individual TIFF files using ImageJ. For each channel, a region containing an enriched signal was selected, and analysis was restricted to this area. A mask was generated to define the region of interest, focusing on the relevant nuclear signal while excluding background. The two channels of interest for this study were the green channel (the “target” channel) and the red channel (the “tester” channel). Colocalization between channels was then evaluated on a pixel-by-pixel basis. Pearson correlation coefficients (PCC) were calculated to quantify the degree of overlap between signals, where higher PCC values indicate stronger colocalization and lower values indicate weaker or no correlation. 10 biological replicates were analyzed using this method to evaluate colocalization across different tissues. PCC values were imported into Prism for statistical analysis across comparisons using a one-way ANOVA + Dunnett’s statistical test.

### RT-qPCR analysis

To measure gene expression, RNA was extracted from S2 cells with RNAeasy Mini kit (Qiagen) and used to generate cDNA with PR1MA qMAX cDNA Synthesis kit (MidSci). 15ul qPCR reactions were carried out with 7.5ul of qMax SYBR green qPCR master mix (MidSci) and 3ul of prepared cDNA. All primer pairs were validated by running serial dilutions and obtaining the R value of >0.9 from the resulting Cq values (see Table S1 for primer sequences). RT-qPCR was performed using the Azure Cielo Real-Time PCR platform. Each sample was run in technical triplicates. Cq values were normalized to expression of beta-tubulin, and the ΔΔCT was calculated for each knockdown and primer set.

### Flow cytometry

S2 cells treated with dsRNA for Nup93-2, Nup93-1, both Nup93-2 and Nup93-1, White and Nup107 were stained with 7-AAD fluorescent dye to interrogate cell viability. Cells were collected at day 2, day 5, and day 7 of the knockdown experiment and washed twice with PBS. Cells were then incubated at 4° C in the dark with 5ul 1x 7-AAD per 1 million cells for 20 minutes. Sample volume was brought to 250ul using PBS before being loaded onto the Flow cytometer. The blue laser, which emits at 640nm, was used to capture 7AAD. Data was gated using FSC, forward scatter and SSC, side scatter to isolate cells of interest. Cell doublets and triplets using FSC-A vs. FSC-NUCLEAR (area v. height) live cells vs. dead cells were gated using forward scatter versus dye fluorescence.

### Western blotting

For tissues, ovaries were dissected from desired CRISPR-tagged flies and homogenized in RIPA lysis buffer. For S2 cells, approximately 2×10^6^ cells were incubated in RIPA buffer for 20 minutes at 4°C, followed by passing through 25g needles. Material was centrifuged for 5 min at max speed, and the supernatants were loaded and ran on 8% gels. Protein was transferred onto PVDF membranes, pre-blocked in PBS+ 5% milk, and blotted with primary antibodies on a shaker at 4°C overnight, then with HRP-conjugated secondary antibodies for 3 hours at RT. Blots were visualized with ECL detection using Analytik Jena ChemStudio Imaging System.

## ACKNOWLEDGEMENTS

We thank members of the Capelson lab for feedback and helpful discussions. We are grateful to Dr. Richard Cripps and Dr. Kim Finley for providing fly stocks and scientific input, and to Dr. Paul Fisher for generously gifting antibodies. We thank Dr. Giacomo Cavalli and members of his lab for scientific discussions, and Dr. Roberto Bonasio and Dr. Emily Shields for scientific input. We also thank Bloomington Drosophila Stock Center and Vienna Drosophila Resource Center for providing fly stocks, Drosophila Genomics Resource Center for providing critical DNA constructs and plasmids, and Developmental Studies Hybridoma Bank for providing antibodies.

## DATA AND RESOURCE AVAILABILITY

RNA-seq datasets have been submitted to GEO (GSE345180) and will be available upon publication (a secure token is available for reviewers). All other relevant data and details of resources can be found within the article and its supplementary information.

**Supplementary Figure S1.**

A. Predicted protein structure comparisons between hNup93 and dNup93 paralogs, derived using AlphaFold and visualized in ChimeraX. Proteins colored by their predicted local distance difference test (pLDDT). Dark blue=very high confidence, light blue=high confidence, yellow=low confidence, orange=very low confidence.
B. Wing imaginal disc phenotypes of Nub-Gal4-driven RNAi against other Nups, as indicated. Note that none of the wing discs show the tumor-like Nup93-2 phenotype. (Confocal microscopy, scale bars for Nup50=200µm, for WT/Nup42=202.8µm, for Nup133/Nup210/Nup62/Mtor=244µm)

**Supplementary Figure S2.**

A. PCA analysis of RNA-seq datasets in Figure 2 by individual replicates from indicated genotypes/conditions.

**Supplementary Figure S3.**

A. RNA-seq-derived normalized expression for CG15059, CG31909 and Ndae1, which are additional genes present in Polycomb domains spanning either Upd1/2/3 or Wg/Wnt genes. (Wald tests adjusted using Benjamin-Hochberg false discovery rate (FDR) analysis performed in R using DESeq2 package. n=3, ns=not significant, *p<0.05, **p<0.01, ***p<0.001).
B. Immunofluorescent staining of *Wingless (Wg)* in control vs Nup93-1 depleted or Nup93-2-depleted larval wing discs (20x objective, 1x zoom, scale bar is 20um). Tissue was stained with DAPI (blue), Lamin (green) and Wg (red).

**Supplementary Figure S4.**

A. RT-qPCR analysis of knockdown levels for RNAi treatment against Nup93-1, Nup93-2, both Nup93-1 and Nup93-2, Nup107 and Ph in S2 cells. Day 7 knockdowns were used in Figure 4A-B.
B. Manual cell counts of indicated RNAi treatments of S2 cells over a 7-day time course.

**Supplementary Figure S5.**

A. Quantification of NPC numbers from the top-of-cell regions (nuclear surfaces, single z slice), using mean grey value of mAb414 via ImageJ.
B. Schematic of the wing disc, indicating the region shown in Figure 5D (created in BioRender). Area of RNAi knockdown indicated by white dashed line.

**Supplementary Figure S6.**

A. Immunofluorescent staining of indicated tissues from HA-Nup93-1 larvae or adult flies, using HA (green) and mAb414 (red) antibodies. DAPI (blue) is included.
B. Immunofluorescent staining of indicated tissues from FLAG-Nup93-2 larvae or adult flies, using FLAG (green) and Nup154 (red) antibodies. DAPI (blue) is included.
C. Western blot validation of the newly generated Nup154 Antibody on protein extracts from S2 cells, treated with control (White) or 2 versions of Nup154-targeting dsRNAs. Tubulin antibodies are used as a loading control.

**Supplementary Figure S7.**

A. Immunofluorescent staining of S2 cells, transfected with Nup93-1-Myc and Nup93-2-GFP constructs, stained with GFP (green) and Myc (red) antibodies and DAPI (blue) (Confocal microscopy, 63x objective, scale bar 4µm).
B. Quantification of co-localization between Nup93-1-Myc and Nup93-2-GFP signals, and between each and mAb414, in S2 cells One-way ANOVA + Dunnett’s statistical analysis in Prism (n=29 cells per condition, 3 biological replicates, ns=not significant, ****p<0.0001).

